# Transferability of Geometric Patterns from Protein Self-Interactions to Protein-Ligand Interactions

**DOI:** 10.1101/2021.10.01.462692

**Authors:** Antoine Koehl, Milind Jagota, Dan D. Erdmann-Pham, Alexander Fung, Yun S. Song

**Affiliations:** Department of Statistics, University of California, Berkeley, CA 94720, USA; Department of Electrical Engineering and Computer Sciences, University of California, Berkeley, CA 94720, USA; Department of Mathematics University of California, Berkeley, CA 94720, USA; Chan Zuckerberg Biohub, San Francisco, CA 94158

**Keywords:** Transfer Learning, Structural Biology, Ligand Docking

## Abstract

There is significant interest in developing machine learning methods to model protein-ligand interactions but a scarcity of experimentally resolved protein-ligand structures to learn from. Protein self-contacts are a much larger source of structural data that could be leveraged, but currently it is not well understood how this data source differs from the target domain. Here, we characterize the 3D geometric patterns of protein self-contacts as probability distributions. We then present a flexible statistical framework to assess the transferability of these patterns to protein-ligand contacts. We observe that the level of transferability from protein self-contacts to protein-ligand contacts depends on contact type, with many contact types exhibiting high transferability. We then demonstrate the potential of leveraging information from these geometric patterns to aid in ligand pose-selection problems in protein-ligand docking. We publicly release our extracted data on geometric interaction patterns to enable further exploration of this problem.

## 1. Introduction

The majority of FDA approved drugs are small organic molecules that exert their effects by binding to various proteins. Experimental discovery of novel drug-like compounds is expensive and throughput-limited. As a result, there is significant interest in developing computational approaches to model protein-ligand binding. Classical methods in this area try to approximate the physics of proteins and ligands to predict the energy of specific protein-ligand binding events. Recently, there has also been interest in applying modern machine learning to this problem. While machine learning has produced some successes, deep learning in particular has not yet had nearly the same impact on modeling of protein-ligand binding that it has had on protein folding [1, 2]. A major contributing factor to this disparity is the fact that there are far fewer experimental structures of protein-ligand interactions (~18,000 [3]) compared to structures of individual proteins (~ 180,000 [4]). This discrepancy is amplified by the fact that while protein self-interactions scale linearly with chain length, protein-ligand contacts are restricted by the relatively constant size of the ligand. In this work, we develop statistical analyses to connect the larger dataset of individual protein structures to the problem of modeling protein-ligand binding.

Since the laws of physics are universal across proteins and ligands, an appealing strategy to work around the scarcity of protein-ligand structures is to transfer knowledge from datasets and models of individual protein structures. Correct protein folding relies on the formation of hydrogen bonds and van der Waals interactions, the same kinds of interactions that allow ligands to bind to proteins (Figure 1). Recently, Polizzi and DeGrado [5] used the idea of transferring geometric interaction patterns from those generated by protein folding (protein self-contacts) to those between proteins and ligands (protein-ligand contacts). They defined the *van der Mer* (vdM), a structural unit that represents interaction geometry between an amino acid and a specific interacting chemical group (iCG). They then calculated vdMs for the iCGs found in amino acids by analyzing a large set of individual protein structures. Finally, they used these vdMs to design a protein to bind a particular ligand. The key insight in this work was that the iCGs found in amino acids are also common in drug-like molecules, and that the interaction patterns for these iCGs might be similar between protein self-contacts and protein-ligand contacts. However, Polizzi and DeGrado did not present a large-scale evaluation of the latter hypothesis.

**Fig. 1.**
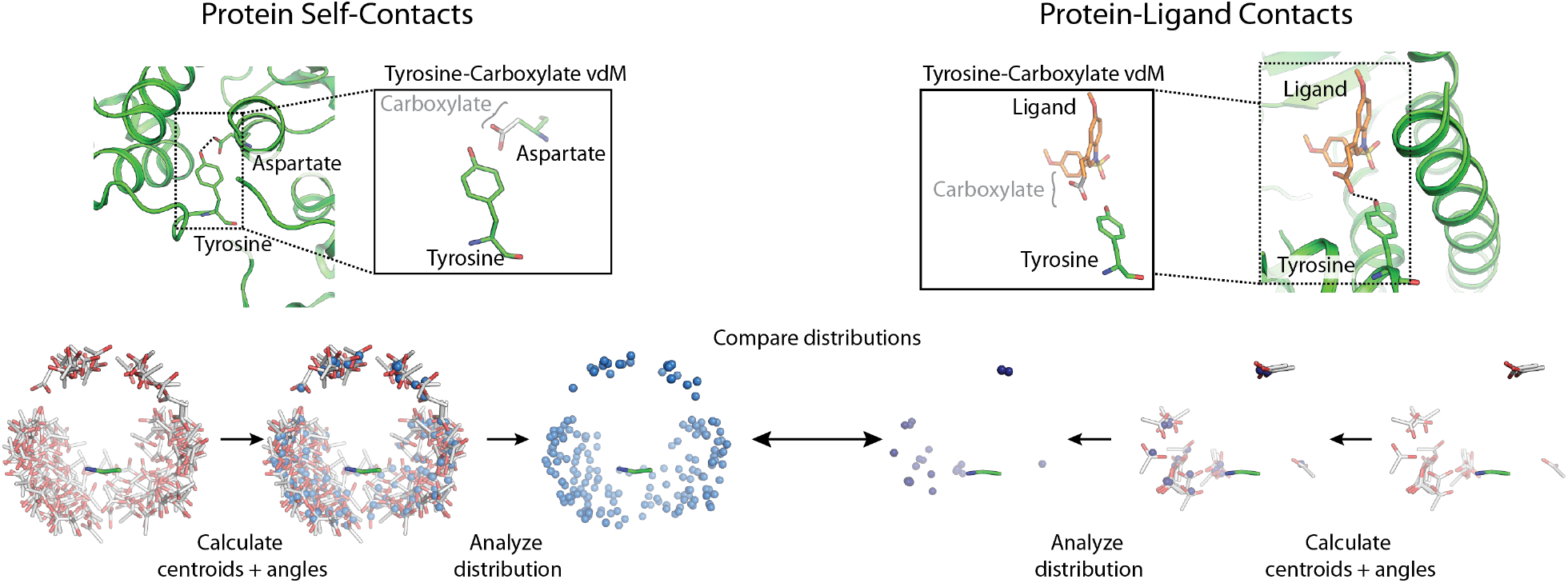
Many important chemical groups in amino acids are also common in drug-like molecules. We seek to measure the transferability of geometric interaction patterns for these chemical groups, between protein self-contacts and protein-ligand contacts. Building on earlier work, we study distributions of chemical group geometry relative to amino acid backbones. We rely on summary statistics of the geometry such as chemical group centroid and atomic angles.

In this work, we present frameworks to more thoroughly assess the transferability of geometric interaction patterns from protein self-contacts to protein-ligand contacts. We first extract the distributions of protein self-contact geometry in an easy-to-use format. We then calculate the same distributions for a dataset of protein-ligand structures and devise a statistical test to quantitatively assess the transferability of geometric patterns from protein self-contacts to protein-ligand contacts. We find that the degree of transferability depends on contact type, with many instances of high transferability. We apply our findings to the important problem of ligand docking and demonstrate the potential of leveraging these transferable contact distributions to improve ranking of candidate ligand docking poses.

## 2. Related Work

Computational, structure-based models of protein-ligand binding are most widely used in the context of ligand docking [3, 6, 7]. Methods for ligand docking rely on scoring functions to evaluate the number and favorability of interactions between the protein and the ligand, but state-of-the-art scoring functions still do not deliver reliably accurate results [6]. Classical scoring functions try to approximate the physics of protein-ligand interactions in order to score the favorability of a ligand binding pose [8, 9]. Recent work suggests that one can supplement these physics-based scoring functions with pattern-based metrics that attempt to recapitulate favorable interaction modes [10]. In particular, methods that maximize structural similarity with a known ligand binding pose [11], or that ensemble predicted poses of multiple ligands [6, 12] have been shown to lead to significantly better enrichment of correctly ranked compounds and poses. However, these methods still rely on either an experimental structure of the binding site with a ligand, or the prior knowledge of multiple ligands that target a particular site, restricting their utility to well-studied protein systems.

There has also been significant interest in using deep learning to learn how to score proteinligand interactions based on 3D structure [13–15]. Methods have been developed based on graph neural networks [13], 3D convolutional neural networks [14], and SE(3) equivariant neural networks [15], among other architectures. However, these models often struggle to maintain accuracy under even slight domain shifts, likely because the available experimental protein-ligand structures are small in number and contain biases that are not well understood [3, 16, 17]. Machine learning methods that have been successful in this space have used very carefully constructed datasets and relatively low capacity models [7].

Deep learning methods have had significant impact in the related problems of protein folding [1, 2] and ligand-based antibiotic discovery [18], where much larger datasets are available. There has been previous work exploring transfer from small-molecule only data to modeling of protein-ligand interactions [19, 20]. Transfer of patterns from individual protein structures to protein-ligand contacts has also been implemented in classical scoring functions at the level of single atoms or bonds [21]. We seek to explore transfer for significantly larger groups of atoms. There has also been previous work in extracting geometric patterns of protein self-contacts at the scale we examine [5, 22]. However, these extracted patterns are either not based on a modern set of experimentally resolved protein structures or do not allow easy measurement of transferability to protein-ligand contacts. To our knowledge, our work is the first to provide precise estimates of the distribution of protein self-contact geometry and the first to examine how these distributions transfer to a collection of protein-ligand structures.

## 3. Methods

### 3.1. Datasets

Protein self-contacts were extracted from the same subset of the Protein Data Bank (PDB) described by Polizzi and DeGrado [5]. In brief, their selection criteria involved resolution (*d*_min_ ≤ 2.0 Å), refinement quality (*R*_obs_ ≤ 0.3), geometric fidelity (Molprobity [23] score ≤ 2.0) and nonredundancy (chains with greater than 30% sequence identity were discarded). This resulted in a database of 9189 unique chains from 8734 PDB structures.

Native protein-ligand contacts were extracted from a combination of the “DCOID” dataset [7] and the 2020 version of the core set of PDBBind [3]. In order to reduce redundancy across this dataset, for all protein-ligand structures with the same UniProt Accession ID, we first clustered structures with ligand Tanimoto similarity ≥ 0.5, and within clusters, removed structures with identical sets of binding pocket residues that contributed to hydrogen bonding contacts. These criteria were established to maximize data retention while also removing obvious duplicate instances of a particular ligand-binding pocket. This yielded a final test set of 5,926 protein-ligand structures. Lists of PDB files for protein self-contacts and protein-ligand complexes are provided in our public dataset release.

Finally, to apply our method to protein-ligand docking, we used the dataset of docking poses for various protein-ligand complexes generated by Paggi *et al*. [6]. They used Glide [8], a widely used docking method, to generate candidate poses for a variety of protein targets of pharmacological interest, each with a number of different ligands.

### 3.2. Contact extraction

As in the work by Polizzi and DeGrado, we extracted data for 20 iCGs which are present in amino acids and therefore well represented in protein self-interactions. These are listed in Supplemental Table S1. To extract protein self-interactions from the PDB data, we first protonated the structures using REDUCE [23] and then identified contacts using PROBE [24]. Flexible regions of a structure were ignored during the contact extraction process by setting maximum crystallographic B-factor and minimum atomic occupancy thresholds of 40 Å^2^ and 0.99, respectively. We extracted interactions that were labeled by PROBE as either hydrogen bonds or close van der Waals contacts. Each interaction was initially labeled as between two different residues in the protein that are separated by at least 7 amino acids to avoid bias from proximity effects. We picked one residue to serve as the source of iCGs and the other to provide coordinates for the amino acid, and then repeated with the choice flipped. After choosing a residue to serve as the source of iCGs, we identified all iCGs in the residue. We then extracted an observed interaction between that iCG and the other residue, composed of the atomic coordinates of the three backbone atoms (N, C*_α_*, carbonyl C) of the residue and the non-hydrogen atoms in the iCG. The interactions were labeled by whether or not they contained a hydrogen bond and by whether or not any atoms of the amino acid side chain were involved in the interaction.

To extract protein-ligand interactions from the protein-ligand test set, we first preprocessed structures using the Protein Preparation Wizard in Schrödinger [25] instead of REDUCE, which we found to be insufficiently accurate for many of the ligands of interest. We then identified protein-ligand interactions using PROBE. The ligands were then decomposed into strictly non-overlapping iCGs via the Schrödinger Python API using SMARTS style strings listed in Supplemental Table S1. For each interaction between ligand and protein involving an iCG, we extracted atomic coordinates for the iCG and the involved amino acid, and labeled them by whether they contained hydrogen bonding and side chain interactions.

For the docking data from Paggi *et al*. [6], processing was performed identically, except that protein-ligand contacts for each docked pose were identified using Schrödinger’s *poseviewer.py* script. Inspection of numbers of extracted contacts between PROBE and *poseviewer.py* on the DCOID dataset showed almost identical identification of protein-ligand contacts, though some minor discrepancies were observed in the treatment of hydrogen bonds involving thiol

### 3.3. Representing contact geometry

Our representations for contact geometry build on the ideas presented by Polizzi and DeGrado [5] (Figure 1). After extraction from our datasets, each contact contained 3D coordinates for an amino acid and 3D coordinates for an iCG. We sorted these contacts into bins corresponding to the amino acid and iCG pair (400 total bins, from 20 amino acids and 20 iCGs – Supplemental Figures S1 and S2). We then calculated two summary statistics for each contact. First, we looked at the distance between centroids (unweighted center of mass) of the iCG and amino acid backbone. Second, for each atom in the iCG, we computed a statistic from −1 to +1 measuring whether the atom points towards or away from the amino acid backbone (described in detail in Section 4.1). While these statistics do not fully parameterize the 3D geometry of a contact, they have the advantages of not requiring alignment and being robust to noise. In particular, Polizzi and DeGrado noted that aligning contacts on backbone atoms leads to the “lever-arm effect”, through which variance in experimental measurements of the backbone atoms is shifted to the iCG atoms and amplified (Supplemental Figure S3). The statistics we have chosen are invariant to the type of variance that causes the lever-arm effect. We also analyzed contacts after alignment on amino acid backbone in Section 4.3 and part of Section 4.1, but the results of these analyses are robust to the lever-arm effect. For ease of visualization and interpretation, we focus on hydrogen bonding interactions in Section 4. This narrows our scope to the 15 iCGs capable of forming hydrogen bonds. However, we provide equivalent preprocessed data for other close van der Waals interactions in our dataset release.

## 4. Results

### 4.1. Protein self-contacts exhibit clear geometric clustering

Here, we explore patterns in the protein self-interaction data, and begin with the distribution of iCG centroid after alignment on amino acid backbone. For the most part, contacts involving a hydrogen bond showed clear geometric patterns with well-separated interaction modes. This agrees nicely with the power law behavior that Polizzi and DeGrado [5] observed in the sizes of their vdM clusters. By further stratifying our analysis into discrete distance radial bins from the reference backbone's centroid, we uncover a strong underlying distance dependence in the overall distribution. In particular, many amino acids share a common short-distance interaction mode that likely reflects chemical group interactions that involve backbone atoms (blue radial bin, Figure 2). The more distant interactions (green and red bins, Figure 2) likely reflect broad geometric constraints on hydrogen bonding conditioned on the rotameric state of the reference amino acid. Rather than explicitly modeling these distributions, aligning on backbone atoms implicitly takes this into account.

**Fig. 2.**
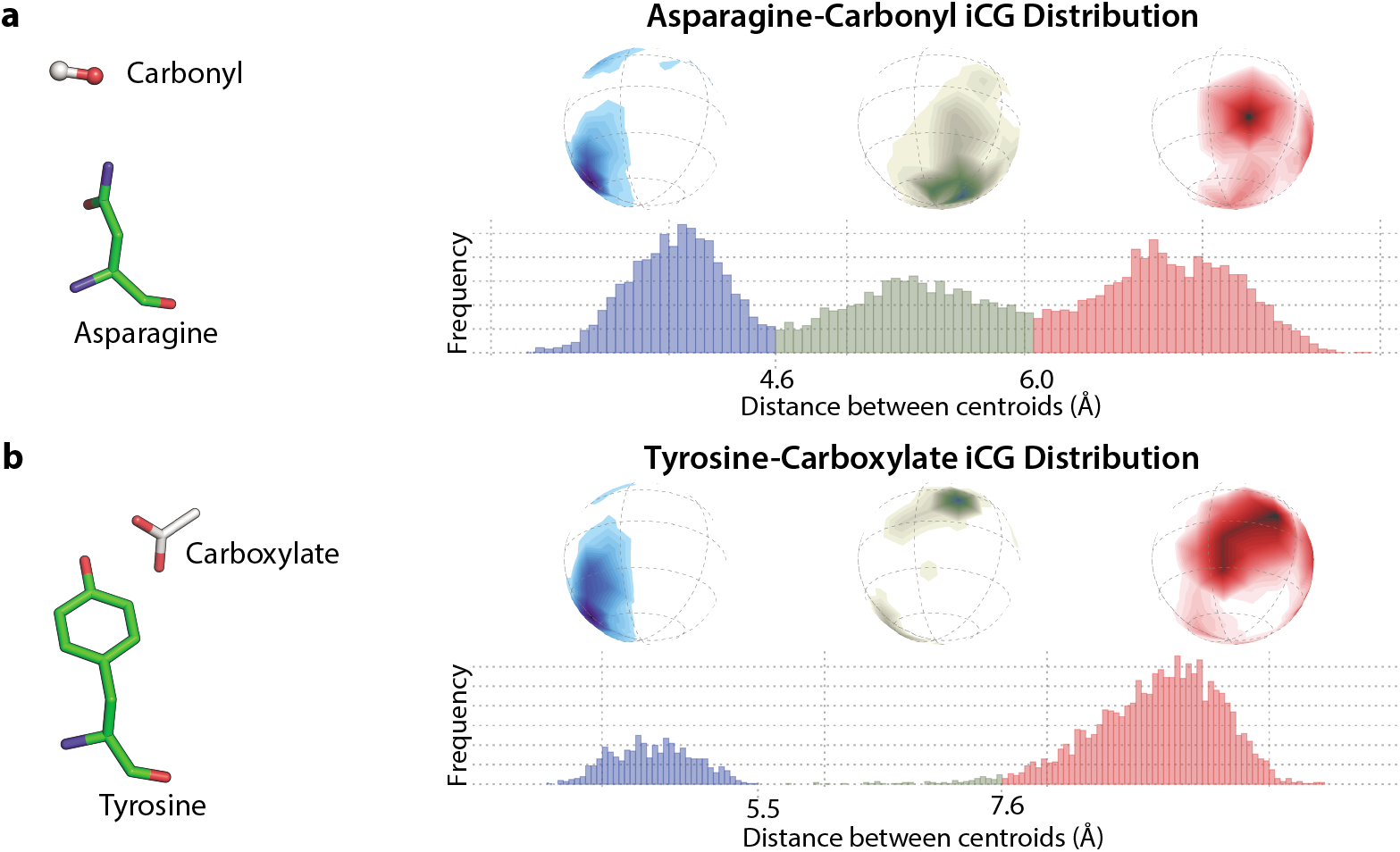
iCGs from protein self-contacts show clear geometric clustering. 3D plot and histogram of iCG centroid density relative to the reference amino acid backbone in different radial bins for two different contact types, Asparagine-Carbonyl (**a**) and Tyrosine-Carboxylate (**b**).

We also explored the distributions of the orientation statistics introduced in Section 3.3. For each atom in the iCG, we computed the vector from the iCG centroid to the atom and the vector from the amino acid backbone centroid to the iCG centroid, and took the cosine of the angle between these vectors (Figure 3a). This value ranges from −1 to +1 for each atom, with −1 indicating that the atom points at the amino acid backbone and +1 indicating that it points away from the amino acid backbone. We observe that the atoms of an iCG that can form hydrogen bonds almost ubiquitously point towards the centroid of the reference amino acid (Supplemental Figure 3a). This is most obvious for smaller iCGs with a single hydrogen bonding atom (Figure 3b), but is also maintained in the indole heterocycle with the nitrogen preferentially oriented towards the reference backbone, and the other atoms (except for the diametrically opposite carbon) distributed almost uniformly elsewhere (Figure 3c). We believe this parameterization of orientation to be more informative than an absolute rotation of the full iCG in 3D, as it provides a direct readout of optimal rotational patterns that maximize hydrogen bonding strength in a contact type.

**Fig. 3.**
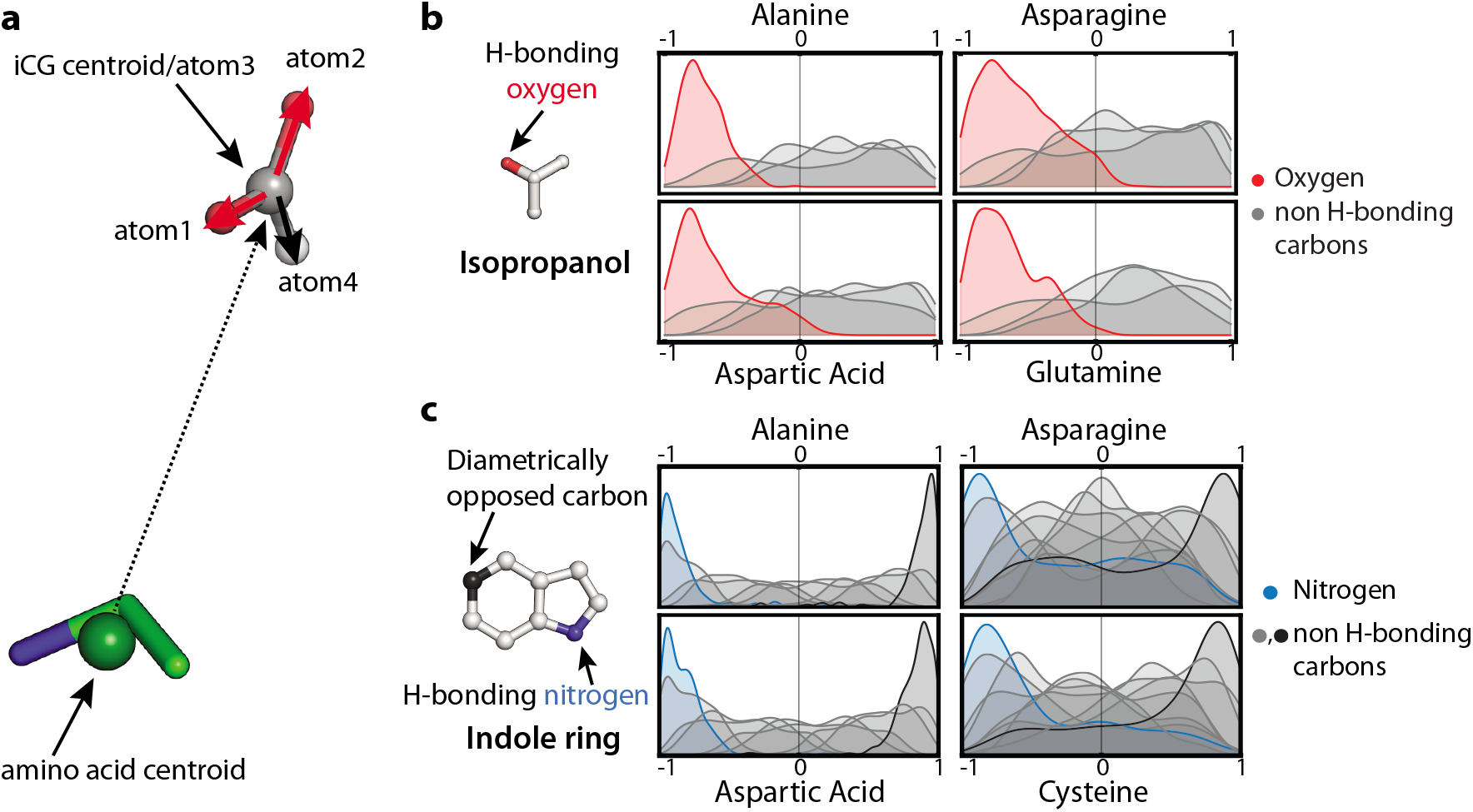
iCGs preferentially orient their hydrogen bonding atoms towards the reference amino acid. (**a**) The atomic angle statistic for a particular atom is the cosine of the angle between the centroid-centroid vector (dotted) and each of the centroid-atom vectors (solid) within the iCG. H-bonding atoms of the iCG are shown in red, while non H-bonding carbons are colored gray. Values range from − 1 (pointing directly to the backbone centroid) to +1 (pointing opposite to the backbone centroid). Orientation distribution of each atom in isopropanol (**b**) and indole ring (**c**) iCGs relative to various reference amino acids. Preferential orientation of the hydrogen bonding atom in each iCG is preserved regardless of whether or not the reference amino acid’s side chain can form hydrogen bonds.

### 4.2. Many geometric patterns transfer to protein-ligand contacts

We next sought to measure how well interaction patterns transfer between protein-protein and protein-ligand instances. It is not immediately obvious what specific notions of “transferability” are useful and feasible to measure. One option would be to apply a statistical test of whether two distributions are identical for each contact type between the protein self-interaction data and the protein-ligand data, using a summary statistic such as the iCG centroid. However, such a test may be too strict, since there are known sources of distributional shift between the two data sources. In their work, Polizzi and DeGrado designed protein-ligand interactions under the assumption that protein-ligand interaction geometries should be near modes in the distribution of protein self-interaction geometries [5]. We chose to use a statistical test that targets a similar use case, by testing for whether protein-ligand contacts have high density under the distribution of protein self-contact geometries.

Concretely, if the true density *f* of a statistic *X* ~ *f* of protein self-interactions (e.g., distance between centroids or angles as described above) were known, a natural approach to identify whether corresponding *K* measurements 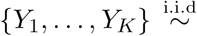 of the protein-ligand interactions fall into high-density regions (according to *f*) would be to investigate the sample mean 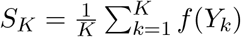 of associated densities. Under the null assumption of *g* = *f* and given sufficiently large *K, S_K_* behaves approximately normal with mean 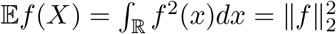 and variance 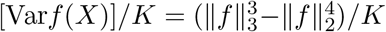, and hence can be converted to *z*-scores to perform one-sided hypothesis testing. In practical settings, *f* is not directly accessible and requires estimation instead. Various standard approaches exist for attaining such estimates, of which a computationally and theoretically particularly appealing one consists of discretizing *X* on a resolution commensurate with the available sample sizes (in our analysis, we guarantee on average ≈ 300 observations per bin), and computing null distributions based on this discretized version. The *p*-values we report are based on this discretized test.

We first applied this test to the distribution of distances between iCG centroids and amino acid backbone centroids for every contact type. This statistic is one-dimensional and able to tolerate the very small sample sizes in the protein-ligand data (Supplemental Figure S2) better than any higher-dimensional statistic. Of all the contact types, 87 had sufficient data for the asymptotic assumptions of the test to be valid (at least 30 samples in protein-ligand data and at least 600 in protein self-interaction data). Of these 87 contact types, 57 had a *p*-value under 0.05 (not transferable) while the remaining contact types had fairly uniform *p*-values (Supplemental Figure S5, left panel) that were not correlated with sample sizes in the proteinligand data (Supplemental Figure S5, right panel). These results indicate that interaction geometries often, but not always, transfer between the two domains. We highlight the full distributions for four contact types that were particularly well sampled, two of which were rejected and two of which were not rejected (Figure 4). In particular, distributions with similar mode structures, but dissimilar density in each mode were rejected by our test as being out-of-distribution (Figure 4, top). This is a potential limitation of our definition of “transferability” depending on the application of interest. We therefore release our preprocessed data and provide density distributions for all contact types in Supplemental Figure S6, so that readers may make decisions about transferability that are appropriate for their own applications.

**Fig. 4.**
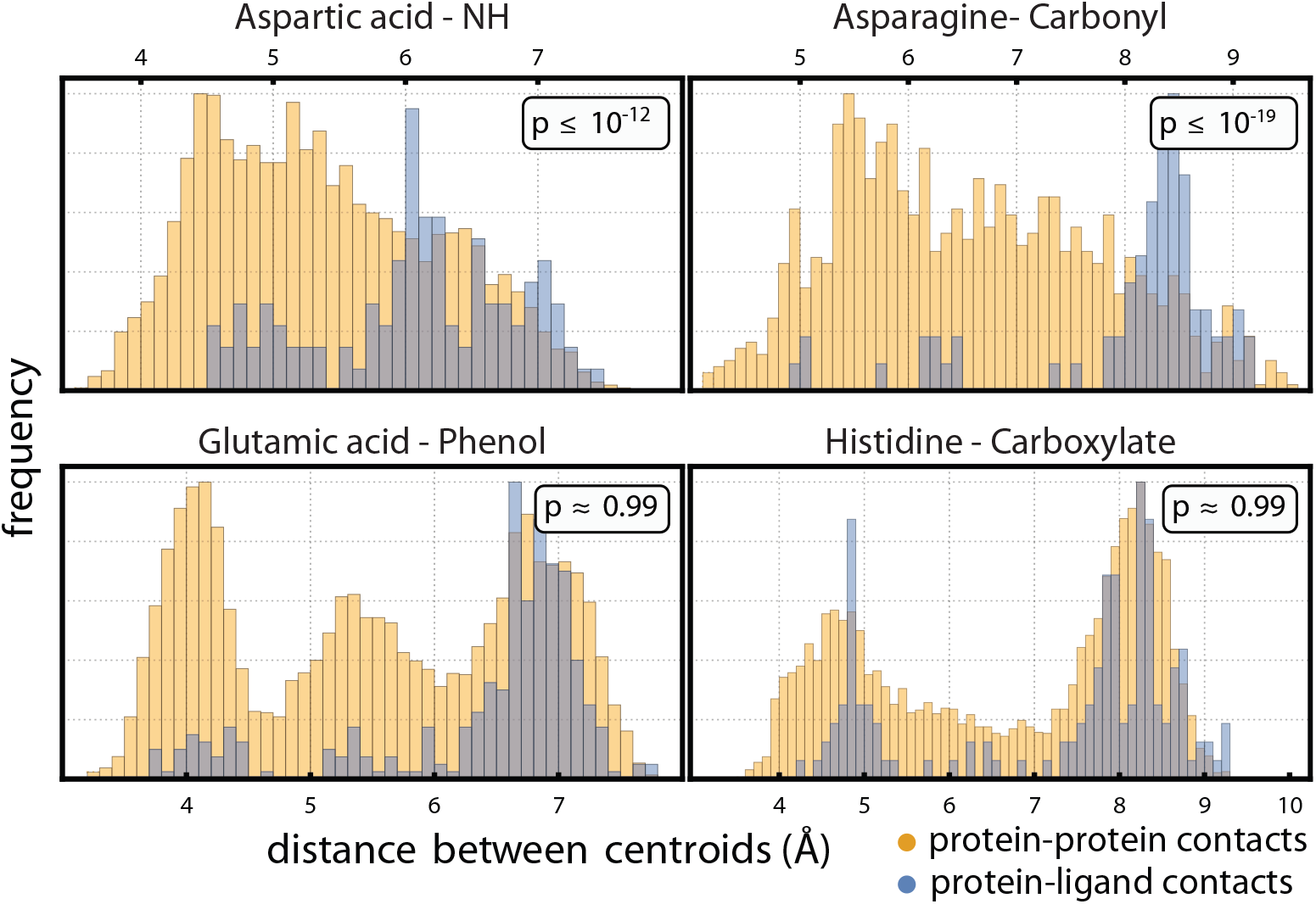
Comparison of protein-protein and protein-ligand backbone-iCG centroid distances for various contact types. Top Row: two cases rejected by our statistical test. The overall mode structures were similar, but with uneven peak densities. Bottom Row: two cases that show similar distributions.

We further used this test to examine how well the iCG orientations defined in Section 3.3 transfer between domains (Supplemental Figure S7). We highlight 4 well-sampled examples, of which two were rejected (Figure 5, top row) and two were accepted (Figure 5, bottom row) by our test. As with the distance distributions, in all cases, the distribution of iCG orientations from protein-ligand complexes fell nicely within the support of those from protein-ligand contacts. Notably, the contact types accepted by our test showed very clear distributional overlaps for all H-bonding atoms, including those not primarily oriented towards the reference amino acid. The rejected samples featured uneven mass distributions relative to the empirical protein-ligand modes; in the cases illustrated here, this is likely a consequence of a contact involving a long and flexible amino acid (Figure 5, top left) or a repulsive interaction between negatively charged groups (Figure 5, top right) in both the amino acid and iCG.

**Fig. 5.**
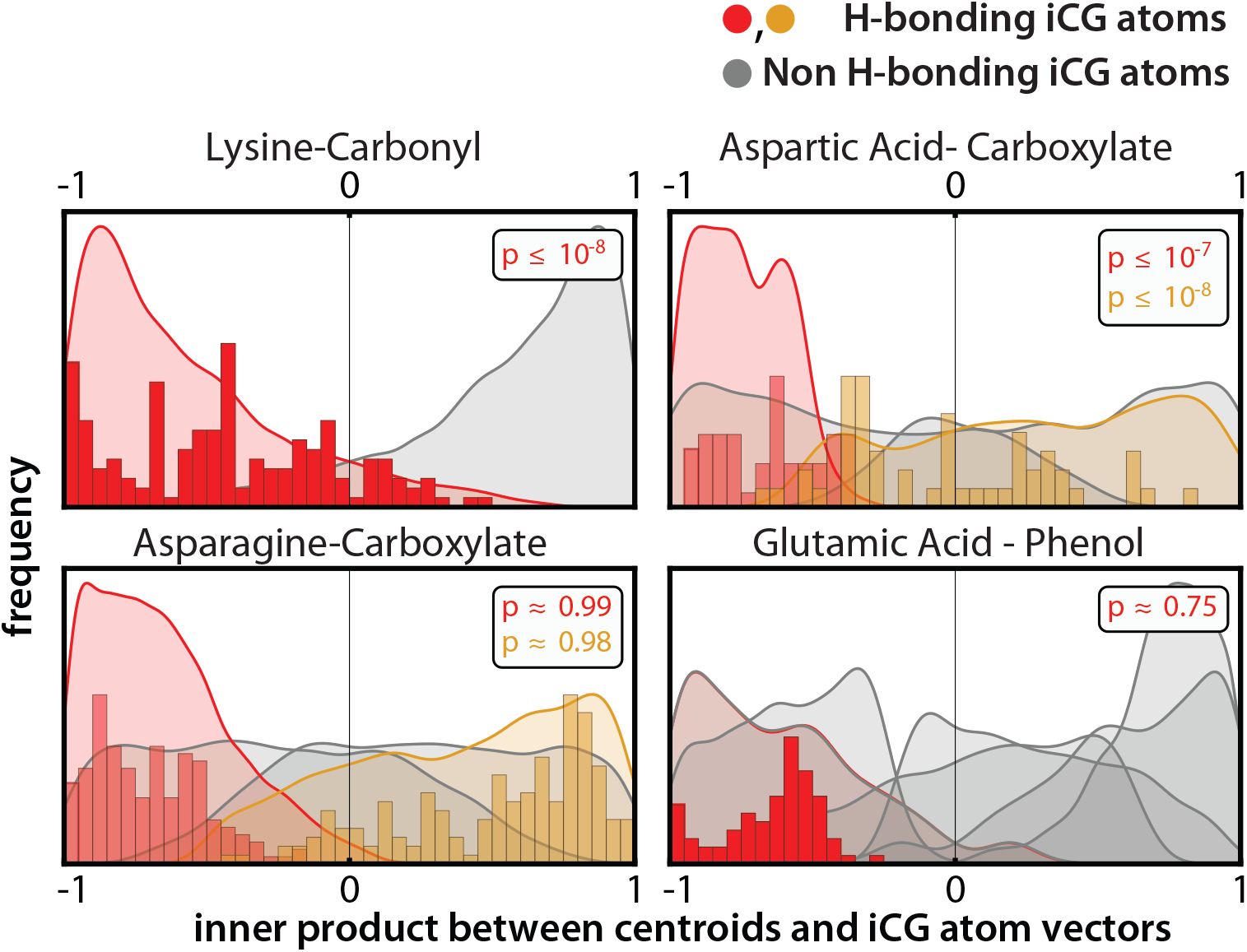
Comparison of protein-protein and protein-ligand iCG orientation distributions for various contact types. Orientations reflect the degree to which a particular atom points in the direction of the reference amino acid’s backbone. Top Row: two contact types rejected by our statistical test. Bottom Row: two contact types that display similar atomic orientations in protein-protein and protein-ligand contacts. Smooth curves represent distributions of atomic orientations from protein self-interaction data; histograms show observations from the protein-ligand dataset.

### 4.3. Application to protein-ligand docking

We explored transferring distributions of protein self-contact geometry to protein-ligand docking. We used the docking dataset described by Paggi *et al*. [6], which includes docking poses for several hundred protein-ligand pairs across 30 different protein targets which are all of pharmacological interest. To apply 3D contact geometry to protein-ligand docking, we constructed a simple method to assign a scalar score to a contact. We only scored contact types with evidence of good transfer to protein-ligand data. Specifically, only contact types with at least 30 samples in the protein-ligand data, distance *p*-value greater than 0.05, and all angle *p*-values greater than 0.05 were used. This left us with 20 contact types. We fit a density for each contact type based on the centroid of the iCG after alignment in the protein self-contact data. To guarantee strictly positive densities, we used a kernel density estimate with a Gaussian kernel on Euclidean distance with bandwidth of 1 A. We then scored contacts from the docking poses by computing their log density under the model for their contact type and then taking the *z*-score of this log density compared to the log densities of protein self-contacts.

We scored all contacts from all docking poses and sorted them by whether they come from an acceptable pose (RMSD to true pose less than 2 A as defined by Paggi *et al*.) or an unacceptable pose (RMSD greater than 2 Å). We also scored contacts from the experimental protein-ligand structures that are available for this dataset. Figure 6 shows the distribution of scores. Note that unacceptable pose contacts are enriched for very negative *z*-scores less than −2.

**Fig. 6.**
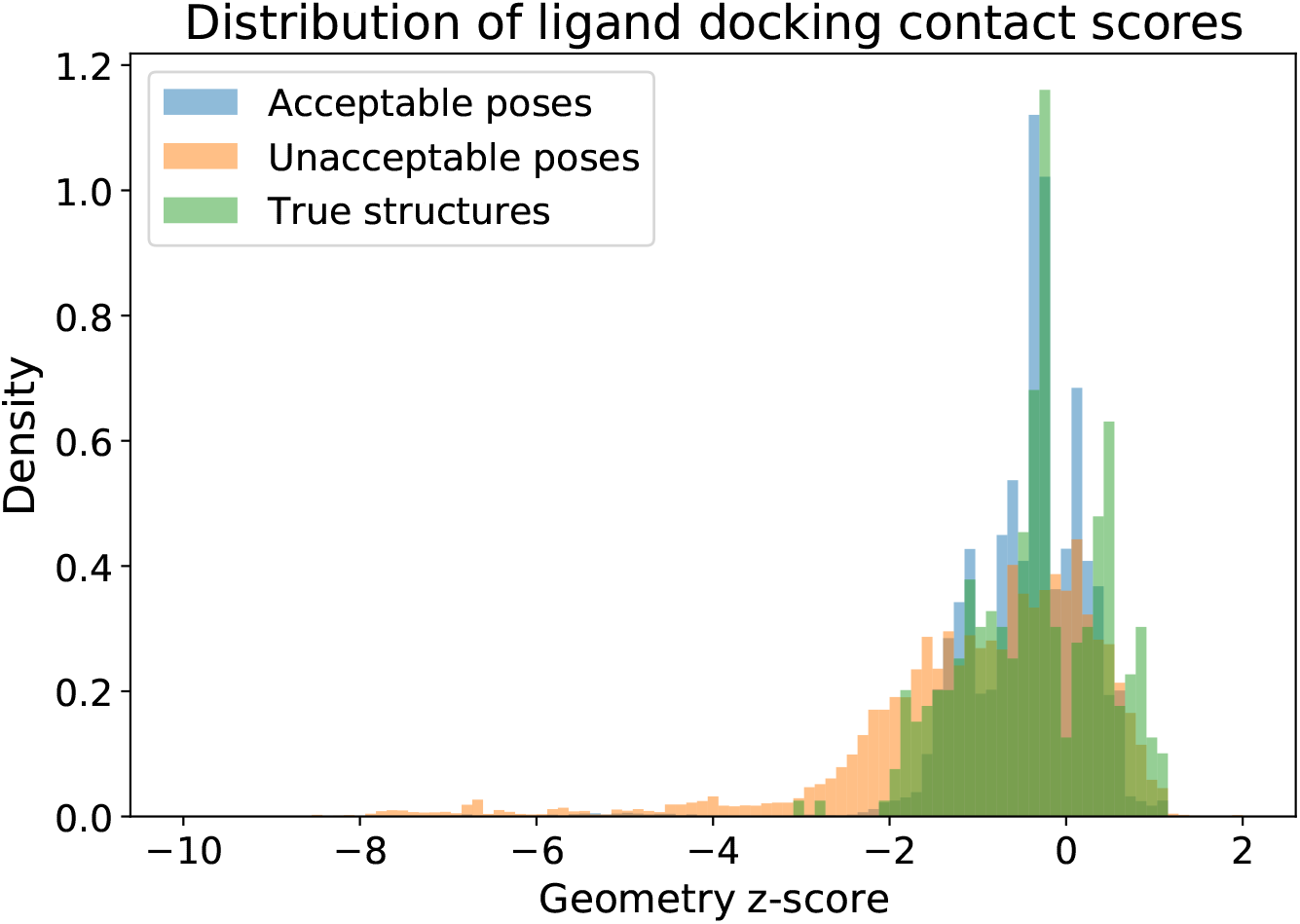
The distributions of protein-ligand contact scores from docking poses is shown for poses of acceptable and unacceptable quality, along with the distribution of true protein-ligand contact scores. These distributions are computed only using contact types with strong evidence of good transferability. Unacceptable pose contacts are enriched for scores less than −2 (16.2% of contacts, compared to 1.4% for acceptable pose contacts and 0.9% for true structure contacts).

These results motivated us to test the effectiveness of simply filtering poses that have a contact with *z*-score less than −2. We examined the top 5 poses of each protein-ligand pair, as ranked by Glide, and found that 836 poses have at least one hydrogen bond of the 20 contact types we examine. Of these, 323 poses were unacceptable and 513 were acceptable. We next calculated the geometric *z*-scores and found that 41 poses had a contact with *z*-score less than −2. Of these 41 poses, 30 were unacceptable and 11 were acceptable, with the unacceptable poses coming from 17 different protein-ligand pairs. These data support the claim that unacceptable poses are enriched by this filter. With the alternative hypothesis that the filter chooses unacceptable poses with higher probability and the null hypothesis that unacceptable and acceptable poses are chosen with equal probability, the *p*-value is 4.5 × 10^−6^. We have thus demonstrated that transferring contact geometry from protein self-contacts can improve protein-ligand docking performance. We anticipate that the increase in performance can likely be improved further by integrating protein self-contact geometry in a more careful manner. This result is also limited by the lack of samples in the protein-ligand data. Collecting more experimental protein-ligand structures will allow the identification of more contact types with good transferability, which in turn will allow scoring a larger number of ligand docking poses. For example, nearly 2,000 poses in this dataset have a contact that could be scored if all contact types were included, compared to 836 in the current analysis.

## 5. Conclusion and Future Work

Our results suggest that while many contact types extracted from protein self-contacts can directly transfer to protein-ligand contact evaluation, the majority show some amount of distributional shift. Qualitatively, many contact types rejected by our test share similar mode structures with the underlying empirical protein self-contact distribution, but with skewed allocation of density into each mode. This observation implies that many amino acids may predominantly employ one of many favorable geometries when interacting with iCGs on ligands. This may reflect a sample size imbalance, or may be the result of optimization of ligand interactions to primarily involve terminal regions of side chains in order to best exploit regions that drive specificity. We therefore provide preprocessed data and introduce a flexible hypothesis testing framework that can be adapted to best suit the application of interest with different levels of stringency. Finally, our preliminary results on the ligand-docking problem indicate the potential impact of applying transferability to drug discovery and other important problems.

## Supplemental Material, Code, and Data Availability

Supplementary Material (with Supplementary Table and Figures), our source code, and preprocessed datasets of protein self-interaction and protein-ligand interactions are available at https://github.com/songlab-cal/contact-geometry.

## Acknowledgments

We thank Nicholas Polizzi and William DeGrado for helpful discussions. This research is supported in part by an NIH grant R35-GM134922. Y.S.S. is a Chan Zuckerberg Biohub Investigator. Antoine Koehl is a fellow at the Miller Institute for Basic Research in Science and is supported by a fellowship from the institute. We acknowledge the UC Berkeley Molecular Graphics and Computation Facility (MGCF) for providing resources and granting access to Schrödinger Software. The MGCF is partially funded by NIH grant S10OD023532.

