## Supplemental Material for "Transferability of Geometric Patterns from Protein Self-Interactions to Protein-Ligand Interactions"

This file contains:

- Supplementary Text
- Supplementary Table S1
- Supplementary Figures S1-S7

[*]These authors contributed equally to this work.

### 1. Definitions of Interacting Chemical Groups

We examine patterns of interaction geometry for 20 different chemical groups that are found in the 20 amino acids. These are the same chemical groups studied by Polizzi and Degrado [1]. We list them here along with the SMARTS string definition for each in Table S1.

### 2. Frequencies of different interaction types

In addition to comparing geometric patterns within each interaction type, one can also ask how common the different chemical groups and amino acids are in protein self-interactions as compared to protein-ligand interactions. We plot both distributions in Figure S1.

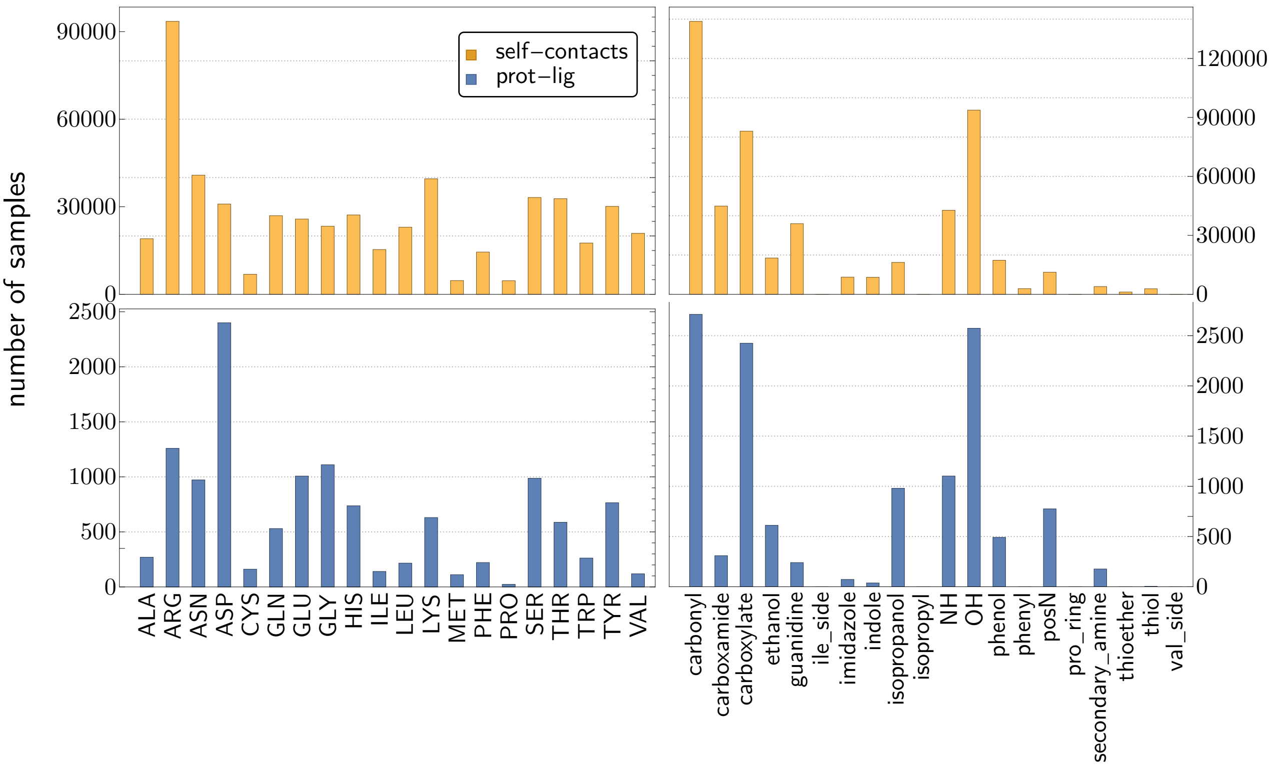


Fig. S1. Frequencies of each iCG and amino acid in both datasets

Table S1. Chemical Functional Group List and Definitions

| **Functional Group** | **SMARTS Pattern** | **Amino Acid** Atoms |
|---|---|---|
| Carbonyl | [C,c]=O | **GLY** C,O |
| Carboxylate | [C,c][CX3](=[OX1])[OH0-,OH] | **ASP** CB, CG, OD1, OD2 |
| Isopropyl | CC(C)[C;!R | **LEU** CB, CG, CD1, CD2 |
| Carboxamide | [C,c][CX3](=[OX1])[NX3H2] | **ASN** CB, CG, OD1, ND2 |
| NH | [C,c][N;H2,H1] | **GLY** CA, N |
| OH | [C,c][O;H1] | **SER** CB, OG |
| Thiol | [CX4;H2][S;H1,H0] | **CYS** CB, SG |
| Thioether | CSC | **MET** CG, SD, CE |
| Guanidine | N=C(N)N | **ARG** NH2, CZ, NE, NH1 |
| Phenol | [O;H1]c1ccccc1 | **TYR** OH, CZ, CE2, CD2, CG, CD1, CE1 |
| N+ | [C,c][N+;H3] | **LYS** CE, NZ |
| Indole | c1c[nH]c2ccccc12 | **TRP** CG, CD1, NE1, CE2, CZ2, CH2, CZ3, CE3, CD2 |
| Imidazole | c1c[n;H0,H1]cn1 | **HIS** CG, CD2, NE2, CE1, ND1 |
| Phenyl | c1ccccc1 | **PHE** CG, CD1, CE1, CZ, CE2, CD2 |
| Isopropanol | [C,c][C;!R]([C;!R])[O;H1] | **THR** CG2, CB, CA, OG1 |
| Ethanol | [C,c][C;H2;!R][O;H1] | **SER** CA, CB, OG |
| Secondary Amine | c[n;H0,H1]c | **HIS** CE1, NE2, CD2 |
| Pro Ring | [$([NX3H,NX4H2+]),$([NX3](C)(C)(C))]1[CX4H]([CH2][CH2][CH2]1) | **PRO** N, CD, CG, CB, CA |
| Val Side | [CHX4]([CH3X4])[CH3X4] | **VAL** CG1, CB, CG2 |
| Ile Side | [CHX4]([CH3X4])[CH2X4][CH3X4] | **ILE** CG2, CB, CG1, CD1 |

### 3. Sample sizes

Sample sizes for the protein-ligand dataset are very small, as shown in Figure S2. We also show sample sizes for the protein self-interaction dataset for comparison.

**Reference Amino Acid**

| iCG | Alanine | Arginine | Asparagine | Aspartic acid | Cysteine | Glutamine | Glutamic acid | Glycine | Histidine | Isoleucine | Leucine | Lysine | Methionine | Phenylalanine | Proline | Serine | Threonine | Tryptophan | Tyrosine | Valine |
|---|---|---|---|---|---|---|---|---|---|---|---|---|---|---|---|---|---|---|---|---|
| Carbonyl | 5794<br>100 | 29845<br>225 | 10893<br>306 | 3123<br>249 | 1347<br>99 | 6895<br>207 | 2482<br>29 | 6523<br>359 | 6683<br>115 | 5836<br>79 | 7528<br>84 | 11261<br>153 | 1239<br>44 | 4118<br>48 | | 7863<br>160 | 8268<br>142 | 4078<br>36 | 6823<br>229 | 8265<br>48 |
| NH | 2496<br>47 | 1220<br>28 | 2514<br>71 | 4146<br>113 | 617<br>16 | 1779<br>28 | 3475<br>82 | 2363<br>350 | 1042<br>50 | 2381<br>8 | 3076<br>39 | 1055<br>14 | 648<br>18 | 2189<br>31 | 1104<br>16 | 2643<br>82 | 2696<br>54 | 1154<br>6 | 2857<br>28 | 3301<br>21 |
| OH | 2565<br>17 | 23204<br>244 | 7413<br>237 | 3474<br>639 | 844<br>3 | 4950<br>120 | 2628<br>350 | 3662<br>103 | 5948<br>243 | 1700<br>5 | 2856<br>11 | 8816<br>125 | 651<br>28 | 1748<br>44 | 449<br>4 | 5924<br>89 | 5827<br>68 | 3566<br>135 | 5083<br>93 | 2409<br>16 |
| N+ | 437<br>20 | 168<br>0 | 591<br>36 | 2647<br>218 | 81<br>0 | 377<br>21 | 2414<br>177 | 489<br>83 | 199<br>0 | 282<br>0 | 514<br>14 | 182<br>0 | 98<br>0 | 310<br>13 | 200<br>0 | 666<br>59 | 626<br>69 | 171<br>0 | 470<br>53 | 337<br>12 |
| Thiol | 70<br>0 | 117<br>0 | 142<br>0 | 68<br>0 | 477<br>1 | 78<br>0 | 42<br>0 | 81<br>0 | 169<br>0 | 55<br>0 | 53<br>0 | 49<br>0 | 55<br>0 | 455<br>0 | 12<br>0 | 150<br>0 | 141<br>0 | 213<br>0 | 341<br>2 | 62<br>0 |
| Ethanol | 703<br>4 | 2087<br>30 | 1725<br>51 | 1339<br>192 | 270<br>0 | 1181<br>23 | 933<br>99 | 835<br>12 | 975<br>57 | 462<br>5 | 722<br>2 | 969<br>14 | 167<br>0 | 480<br>29 | 187<br>0 | 1269<br>30 | 1394<br>12 | 901<br>27 | 1159<br>23 | 708<br>1 |
| 2° amine | 101<br>0 | 245<br>0 | 242<br>0 | 262<br>0 | 67<br>0 | 155<br>0 | 193<br>0 | 172<br>0 | 199<br>0 | 73<br>0 | 172<br>0 | 114<br>0 | 59<br>0 | 290<br>0 | 61<br>0 | 378<br>115 | 281<br>0 | 268<br>0 | 497<br>61 | 112<br>0 |
| Thioether | 23<br>0 | 94<br>0 | 179<br>0 | 25<br>0 | 92<br>0 | 127<br>0 | 18<br>0 | 45<br>0 | 70<br>0 | 24<br>0 | 33<br>0 | 38<br>0 | 5<br>0 | 20<br>0 | | 79<br>0 | 103<br>0 | 72<br>0 | 87<br>0 | 35<br>0 |
| Carboxamide | 1814<br>4 | 4666<br>8 | 4747<br>11 | 3107<br>89 | 460<br>2 | 2962<br>7 | 2371<br>22 | 1987<br>62 | 1705<br>7 | 1471<br>4 | 2166<br>20 | 2479<br>11 | 547<br>13 | 1426<br>0 | 684<br>1 | 3002<br>25 | 3161<br>4 | 1597<br>3 | 2794<br>12 | 1750<br>4 |
| Carboxylate | 1904<br>53 | 26185<br>646 | 5931<br>161 | 1018<br>51 | 412<br>30 | 3441<br>90 | 795<br>46 | 3196<br>80 | 6389<br>152 | 1099<br>30 | 1908<br>22 | 11939<br>246 | 371<br>6 | 869<br>20 | 3<br>0 | 5263<br>309 | 4310<br>208 | 2490<br>33 | 4035<br>233 | 1453<br>9 |
| Guanidine | 1476<br>11 | 751<br>1 | 1883<br>3 | 7622<br>110 | 277<br>0 | 1554<br>8 | 7103<br>27 | 1782<br>26 | 734<br>2 | 818<br>4 | 1861<br>7 | 575<br>1 | 311<br>2 | 929<br>3 | 1135<br>1 | 1995<br>23 | 1816<br>4 | 640<br>5 | 1679<br>1 | 1011<br>1 |
| Isopropanol | 611<br>1 | 1917<br>36 | 1532<br>61 | 1033<br>598 | 181<br>0 | 1196<br>16 | 700<br>87 | 763<br>16 | 817<br>58 | 375<br>2 | 545<br>2 | 821<br>32 | 171<br>0 | 365<br>16 | 142<br>0 | 1184<br>24 | 1607<br>11 | 735<br>9 | 981<br>11 | 560<br>0 |
| Imidazole | 242<br>0 | 612<br>2 | 522<br>2 | 1047<br>17 | 122<br>0 | 339<br>1 | 684<br>10 | 394<br>9 | 504<br>2 | 177<br>0 | 319<br>3 | 292<br>0 | 97<br>0 | 390<br>1 | 129<br>0 | 723<br>17 | 640<br>4 | 417<br>1 | 860<br>3 | 244<br>0 |
| Phenol | 494<br>13 | 1798<br>37 | 1595<br>31 | 1265<br>115 | 512<br>8 | 1156<br>9 | 1162<br>76 | 621<br>8 | 1008<br>51 | 371<br>3 | 772<br>8 | 718<br>34 | 180<br>0 | 587<br>16 | 310<br>1 | 1166<br>42 | 1071<br>11 | 770<br>7 | 1365<br>14 | 419<br>7 |
| Indole | 296<br>0 | 434<br>2 | 674<br>2 | 738<br>9 | 379<br>2 | 587<br>0 | 782<br>2 | 375<br>2 | 464<br>0 | 197<br>0 | 457<br>4 | 187<br>0 | 104<br>0 | 285<br>0 | 248<br>0 | 595<br>12 | 598<br>0 | 374<br>0 | 674<br>2 | 206<br>0 |

Fig. S2. Protein-Ligand sample sizes including only contacts with hydrogen bonds. For each contact type, note the number of observations in the protein-protein(top number-yellow) and protein-ligand(bottom number- blue) datasets

### 4. Lever-arm effect

In their work, Polizzi and DeGrado observed the "lever-arm effect", whereby aligning observed contacts by amino acid backbone amplifies noise in the iCG [1]. We illustrate this effect with a simulation in Figure S3. In Figure S3a, we start out with a true contact geometry between a red triangle and a blue square. In Figure S3b, we sample 50 instances of the true geometry plus noise. The noise has both a component that is applied to all verticies independently as well as a random rotation around the centroid that is applied independently to the triangle and square. In Figure S3c, we then align these instances on the triangle. Although the true geometry is still recognizable after adding the noise, aligning on the triangle amplifies the noise on the square so that the true geometry is hard to see.

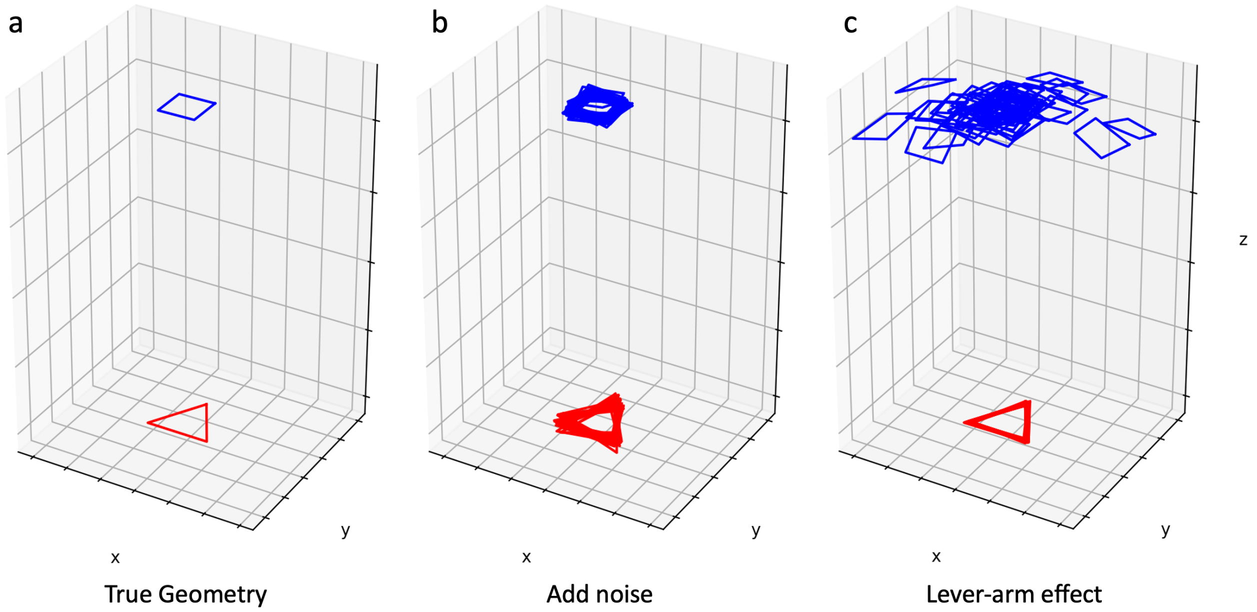


Fig. S3. Simulation of the lever-arm effect. a) The true geometry of a simulated contact between a red triangle and a blue square is shown. b) 50 versions of the contact plus noise are sampled. The noise has a component that is added independently to each vertex as well as random independent rotations of both the triangle and square around their centroids. c) After aligning the noisy contacts on the triangle, the noise of the square becomes amplified so that the true geometry is hard to see.

### 5. Plots of atomic angles

We show the distribution of cosine angles for every iCG atom in every contact type that can have hydrogen bonds in Figure S4. Hydrogen bonding atoms almost always have a strong negative skew indicating that they point inwards towards the amino acid backbone.

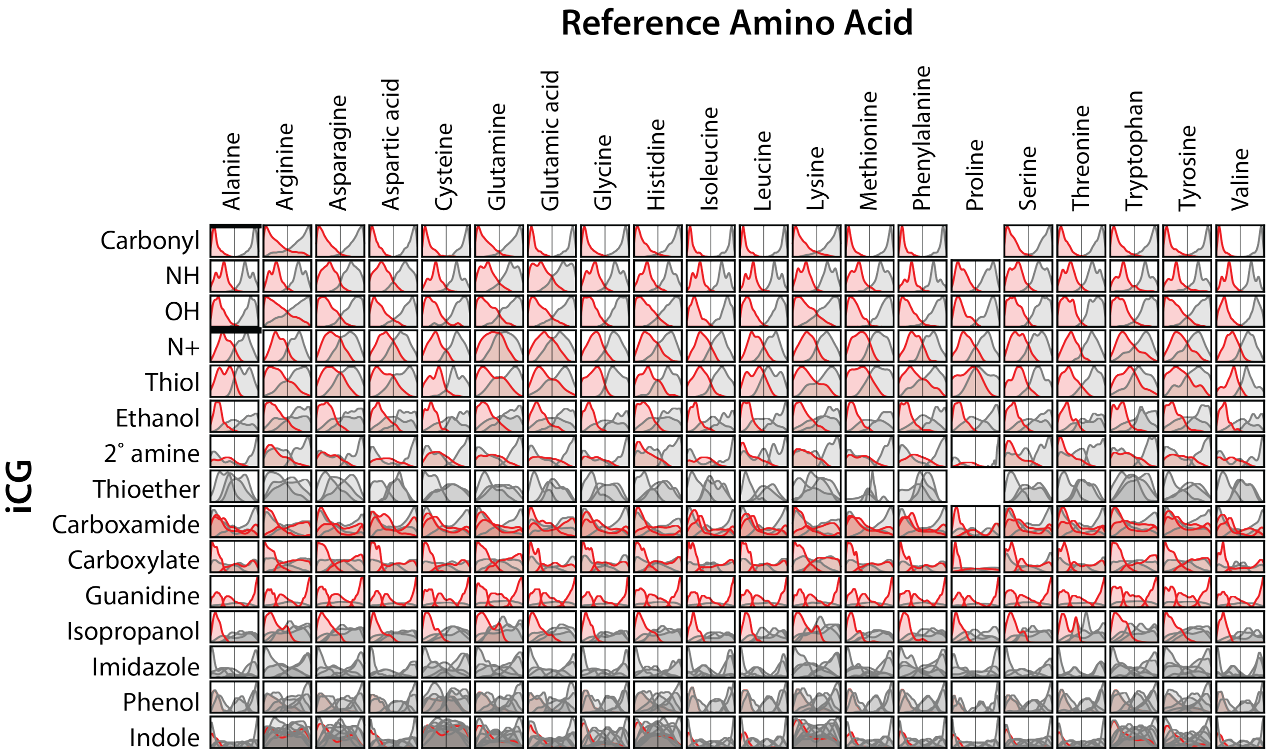


Fig. S4. Orientation of iCG for each contact type relative to the centroid vector. H-bonding atoms of the iCG are shown in red. There is a clear preference for orientations that place H-bonding atoms towards the reference amino acid centroid

### 6. Radius testing $p$-values

For the transferability test that is described in the main text, we had sufficient samples for 87 contact types. The distribution of $p$-values along with the correlation to sample size in the protein-ligand dataset are shown in Figure S5.

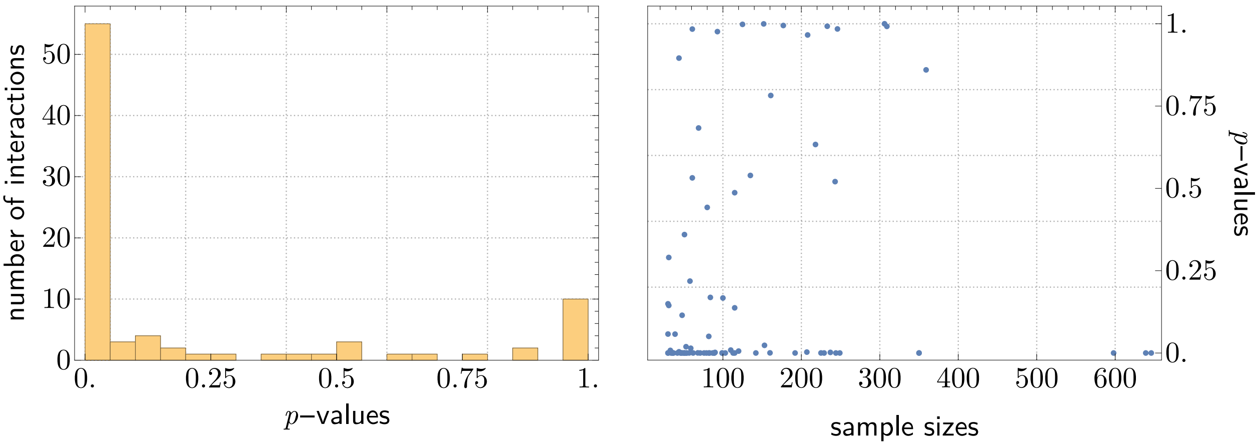


Fig. S5. Distribution of $p$-values for hypothesis test of transferability on the distance between iCG centroid and amino acid backbone centroid. 87 contact types had sufficient samples for the test to be valid and of these, 57 had $p$-values less than 0.05. The $p$-values do not have a clear correlation with sample size in the protein-ligand data indicating that failures to reject are not just due to lack of samples.

### 7. Radius testing distributions

We provide amino acid- iCG centroid distance distributions from protein-protein and protein-ligand contacts. In the majority of cases, both distributions show similar interaction modes - however, imbalances in the density of contacts in each mode will lead to rejection by our test. Figure S6 provides a qualitative view of these distributions to aid in understanding similarities and differences between protein-ligand and protein self-contact contacts.

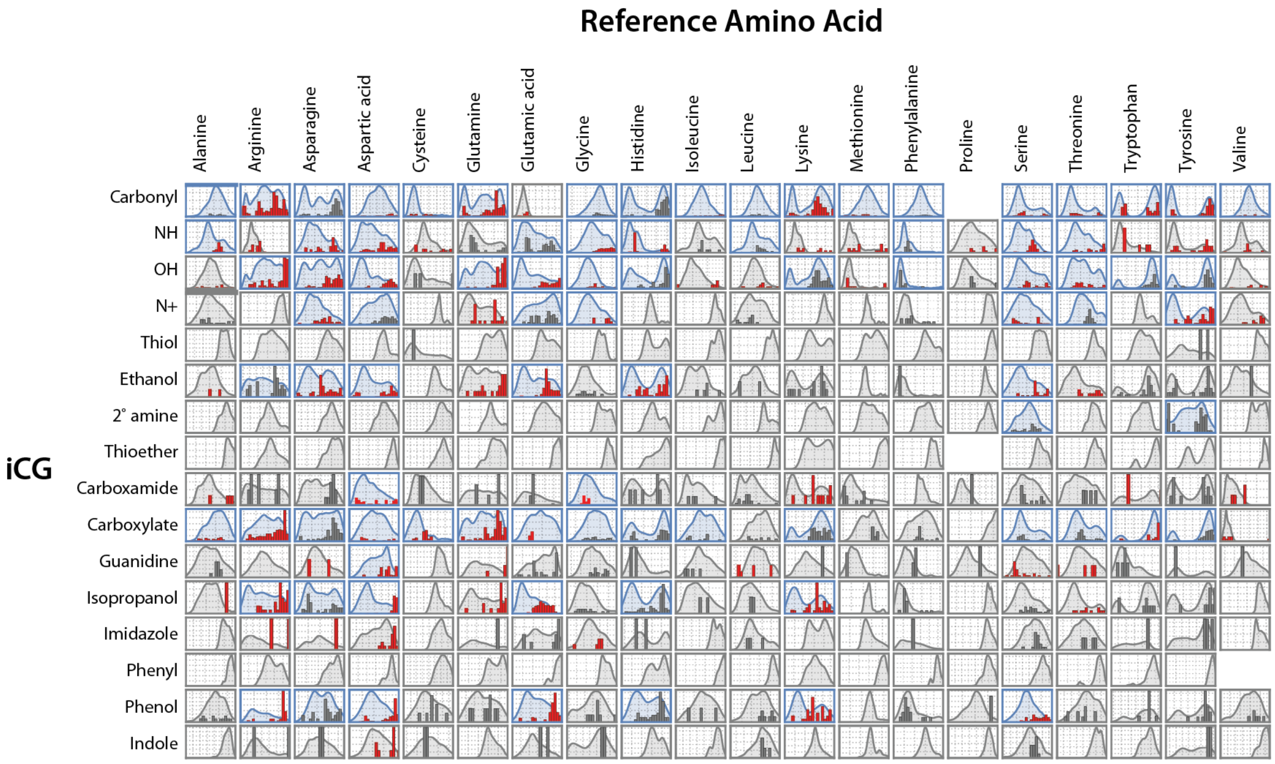


Fig. S6. Distribution of iCG-Backbone centroid distances of all contacts from protein self-interactions (smooth curves) and protein-ligand interactions (histrograms). Contact types that meet the asymptotic assumptions of our test are colored blue, all others are colored grey. Contact types with matching protein-ligand and protein self contact distributions have gray histogram bars, while those rejected by our test are colored red.

### 8. Orientation testing $p$-values

For the orientational transferability test that is described in the main text, we had sufficient samples for 87 contact types. The distribution of $p$-values along with the correlation to sample size in the protein-ligand dataset are shown in Figure S7

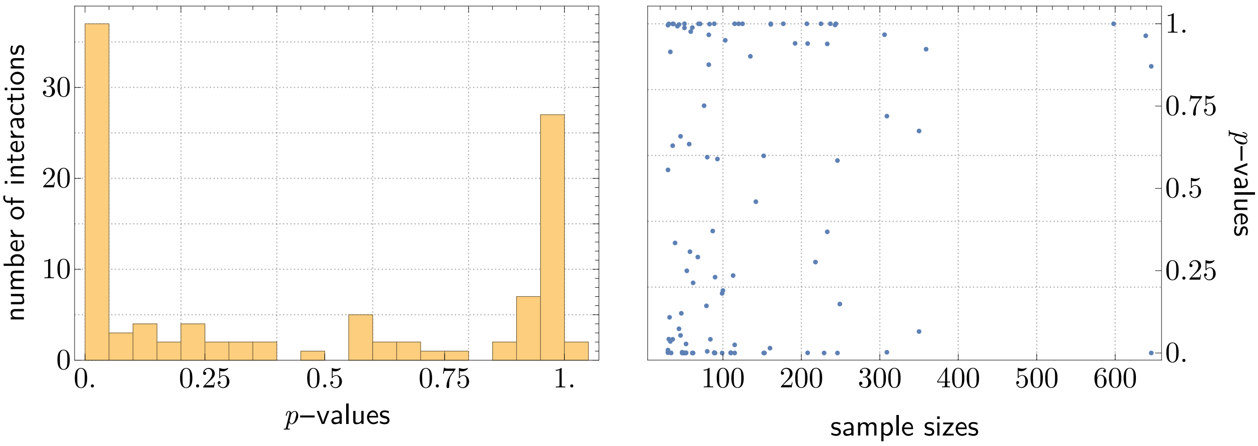


Fig. S7. Distribution of $p$-values for hypothesis test of transferability on the distance between iCG centroid and amino acid backbone centroid. 87 contact types had sufficient samples for the test to be valid and of these, 37 had $p$-values less than 0.05. The $p$-values do not have a clear correlation with sample size in the protein-ligand data indicating that failures to reject are not just due to lack of samples.
